# A probabilistic functional atlas based on extraoperative electrocortical stimulation mapping

**DOI:** 10.1101/2025.09.15.676264

**Authors:** Andrew J. Michalak, Leyao Yu, Amirhossein Khalilian-Gourtani, Alia Seedat, Cassandra Kazl, Christina Morrison, Zachary Resch, Werner Doyle, Peter Rozman, Orrin Devinsky, Patricia Dugan, Daniel Friedman, Adeen Flinker

## Abstract

**Background:** Direct electrocortical stimulation (DES) is the clinical gold standard for identifying eloquent cortex and guiding neurosurgical intervention, yet prior intraoperative DES during awake craniotomies have been limited by intraoperative sampling constraints and density-based rather than probabilistic analyses. The probability of typical and atypical cortical organization has not been fully explored, especially in epilepsy populations. We sought to generate a probabilistic atlas of motor, sensory, and language functions using extraoperative DES in a large cohort of patients with epilepsy.

**Methods:** We retrospectively analyzed 2,124 extraoperative DES trials from 125 patients undergoing intracranial monitoring (2008–2023). Positive and negative trials were mapped to Montreal Neurological Institute space, parcellated with the Human Connectome Project atlas, and analyzed using probability mapping, bootstrapped region-of-interest hit probabilities, hierarchical clustering, and kernel density estimation. Mixed-effects models assessed clinical predictors of language disruption.

**Results:** Probabilistic maps revealed regions of increased likelihood for eliciting functional responses in expected sensorimotor and language territories, but also demonstrated marked variability and deviations from expected cortical locations. Language disruption occurred in 338 trials, motor in 520, and sensory in 370. Instead of observing high probabilities and low inter-patient variability isolated to classic perisylvian locations (e.g., Broca’s and Wernicke’s areas), the likelihood of language disruption followed graded probabilistic gradients with high inter-patient variability. The middle frontal gyrus emerged as a consistent locus of naming and speech arrest. Motor phenomena extended into parietal association cortex. Higher-order experiences, including forced thoughts and feelings of presence, were reproducibly evoked from frontal and temporoparietal sites. Early seizure onset and temporal lobe lesions predicted lower naming disruption probabilities.

**Conclusions:** This extraoperative DES atlas, the largest to date, demonstrates that eloquent cortical functions are organized along probabilistic continua rather than fixed regions. Findings highlight the middle frontal gyrus as a critical language node, extend motor mapping into parietal cortex, and delineate reproducible experiential phenomena. Substantial inter-patient variability underscores the necessity of individualized mapping in surgical planning.

## Introduction

Direct electrocortical stimulation (DES) can identify eloquent cortex and guide safe neurosurgical intervention. By applying current directly to exposed cortex, DES transiently disrupts local neuronal activity and provides causal evidence for cortical sensory, motor, and language functions.[1–6] DES-elicited phenomena typically arise from brain regions serving specific functions, such as motor control in the precentral gyrus and language in perisylvian regions.[7, 8] Although certain regions exhibit a higher probability of eliciting functional responses, there is intra-individual and even greater inter-individual variability, with functional reorganization common with earlylife pathologies. [9] Anatomical lesions, impaired neurocognitive function, and chronic seizures - especially from a young age - can cause atypical localization of cortical representations (i.e. functional reorganization).[1, 10–13]

Studies have tried to assess this variability by deriving generalizable functional atlases from large intraoperative DES cohorts. Some aggregated positive responses across patients to identify cortical representation patterns by analyzing positive trials.[3–6] However, exclusion of negative trials where stimulation elicited no response prevents the assessment of the *probability* of eliciting a response in a region. High density regions are prone to sampling bias and are often more frequently sampled than regions with a lower *a priori* expectation of a functional response. Further, intraoperative DES is limited by cortical exposure, a narrow time window during awake craniotomies, patient cooperation, and electrographic seizures and after-discharges;[14] all could under-represent less classical regions. For instance, the middle frontal gyrus (MFG) has emerged as a critical region for language processing in recent years,[15] yet language disruption in the MFG is under-reported among intraoperative DES studies.[2, 3, 6, 16] Although some studies have computed probabilistic heatmaps and identified regions with a high inter-patient probability and low variability of eliciting a given function[2, 17], it is unclear how accurately these maps represent the true spatial distribution of cortical functional organization.

Extraoperative DES has advantages over intraoperative DES in this context. Patients undergoing presurgical evaluation for epilepsy surgery are implanted with intracranial grids, strips, and/or depth electrodes for several days to weeks.[18, 19] At the bedside, DES can be performed while patients are awake, with fewer time constraints and broader coverage across implanted regions including both resection targets and unaffected cortex, providing positive and negative trials needed to build true probabilistic maps. [11, 20–22]

Using extraoperative DES mapping data from a cohort of 125 epilepsy patients, we addressed limitations of previous intraoperative DES reports by providing probabilistic maps of language, sensory, and motor phenomena. Hierarchical clustering of anatomical regions in Human Connectome Project Multi-Modal Parcellation (HCP-MMP) revealed robust structure–function relationships, including expected motor somatotopy alongside distributed language responses. Importantly, we revealed a continuous probabilistic gradient for speech arrest and naming, with MFG emerging as a critical site for language disruption. Clinical variables related to cortical development and seizure onset influence the likelihood of language disruption. Lastly, we provided evidence from our large cohort on higher-order sensory and experiential phenomena localization, previously limited to small case series. Together, these findings establish an extraoperative DES-based atlas of functional topography, highlight high inter-patient variability in structure-function relationships, and reinforce the need for individualized mapping prior to resection.

## Methods

### Patient selection and participant details

This retrospective observational study included all consecutive patients between March 2008 and January 2023 who gave consent to participate in research at New York University Langone Hospital and who underwent intracranial grid/strips placement and extraoperative DES as part of routine clinical care. Patients were implanted with subdural grids/strips with or without depth electrodes (ADTech Medical Instrument Corp.). We collected demographic and clinical information including age of epilepsy onset, location of seizure onsets, and the presence and type of any lesion. A patient was considered lesional if there was evidence of a lesion associated with their epilepsy (e.g., tumor, vascular malformation, focal cortical dysplasia, gliosis, etc.) based on magnetic resonance imaging (MRI) or post-resection histopathology. Patients with incidental lesions (e.g., small cavernous malformations) unassociated with the seizure network on intracranial EEG were classified as non-lesional.

### Electrical stimulation protocol

Extraoperative DES was performed at bedside following a clinical protocol[18] after antiseizure medications were resumed, as described previously.[23] Briefly, DES was performed using a NicoletOne Cortical Stimulator (constant current output, pulse width 500 microseconds, pulse frequency 50 Hz, maximum train duration 5 seconds). The stimulus was applied to adjacent electrode pairs, starting at 1-2 mA and gradually increased until a functional response or maximum current (15 mA for subdural electrodes or 8 mA for depth electrodes) was reached.

Patients performed motor and sensory tasks, continuous speech (e.g., stating months of the year, days of the week, or a memorized prose paragraph such as The Pledge of Allegiance or a prayer), visual and auditory naming tasks, and auditory comprehension trials (Figure 1A). For each electrode pair, the patient began counting while stimulation was delivered at random intervals, starting with the lowest current level and gradually increasing until a behavioral deficit was observed or threshold stimulation was reached. If a language deficit was observed, the task was repeated with the same threshold stimulation parameters until the same deficit occurred on at least two of the three trials. If there was no symptom or behavioral deficit across any of the language tasks, the target electrode was considered “cleared” (i.e., noneloquent for the language paradigms assessed). If another phenomenon was elicited (e.g., significantly distracting or debilitating motor/sensory phenomena or epileptic auras), stimulation was terminated for that pair. If seizures or afterdischarges occurred, the clinician could administer additional antiseizure medication to reduce cortical excitability and allow testing to continue. Seizures or epileptic auras elicited by stimulation excluded the electrode pair from further analysis. If afterdischarges occurred with subthreshold current, the electrode pair was excluded from further analysis. However, if afterdischarges occurred at the threshold current but no symptoms were elicited, the electrode pair was considered “cleared.” Since bipolar stimulation is performed pairwise across neighboring electrodes, both electrodes were labeled for each trial. We use the terms positive and negative “trial” to indicate the presence or absence of a behavioral response during stimulation of an electrode pair. In contrast, “hit” and “clear” refer to individual electrodes that were part of a positive or negative trial, respectively. Thus, the total number of electrodes represented in figures is twice the number of trials since a positive or negative trial would yield two hits or clears.

**Fig. 1.**
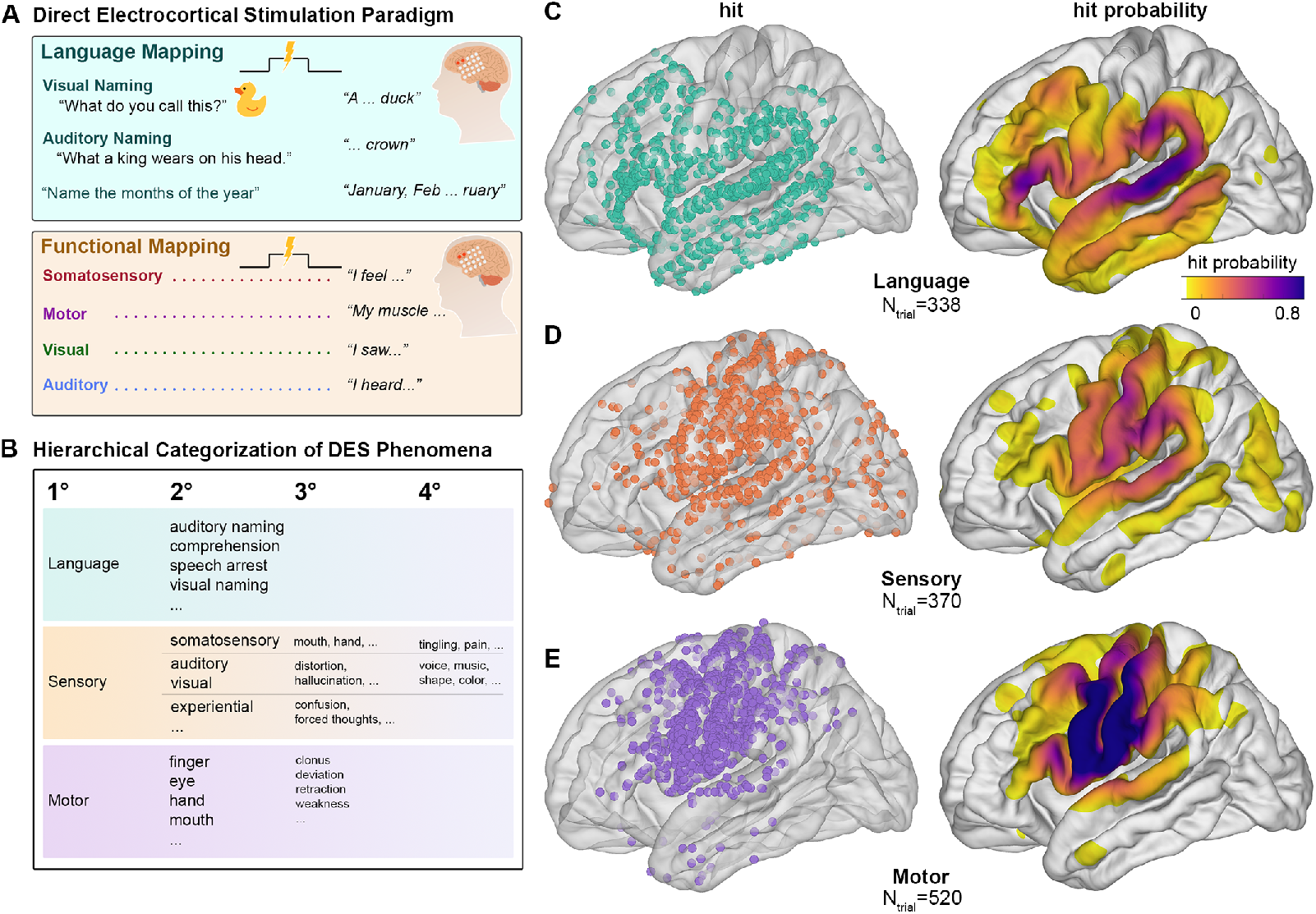
Task paradigm and hit distribution on template brain. (A) Experimental paradigm depicting the tasks during DES. (B) Hierarchical classification of the phenomena. (C) Hit and hit density of primary categories.

Elicited symptoms were categorized using a hierarchical taxonomy structured as “modality → body part → phenomenon” (Figure 1B). For example, a motor hit could be recorded as “right hand clonic” or a sensory phenomenon such as “left hand and arm tingling.” Sensory phenomena were categorized as somatosensory (e.g., tactile), visceral, visual, auditory, or experiential (e.g., déja vu or autoscopy). Visual and auditory hallucinations were further categorized as simple (e.g., a flash of light or beeping sound) or complex (e.g., a human face or hearing formed words). Language mapping results were subdivided into speech arrest, auditory or visual naming, and comprehension. A full table of recorded phenomenology is available in Supplementary Table 1.

### Anatomic localization

Visualization and statistical analyses were done by probability density plots and by regions of interest (ROIs). For each patient, the electrodes were localized onto the T1-weighted MRI in patient space then linearly transformed into Montreal Neurological Institute (MNI) 152 space for group analysis. Next, the cortical parcellation was derived from T1-weighted MPRAGE images using the FreeSurfer pipeline.[24] The Desikan-Killiany atlas was used for analysis of gross gyral anatomy, and for analysis of functional regions of interest (ROIs) we used the Glasser Human Connectome Project Multi-Modal Parcellation Atlas (HCP-MMP), an atlas used in functional and structural connectivity research containing cortical segmentations relevant to cognition, perception, and disease.[25, 26]

### Statistical analysis

All statistical analysis and visualization were performed in MATLAB (Natick, MA) and Python 3.10. [27, 28] Data points across patients were discretized on the cortical surface using MNI coordinates to create surface probability maps for language, motor, and sensory findings. To explore gross cortical representations, we aggregated all patient trials to generate probability maps (positive stimulation trials / (positive stimulation trials + negative stimulation trials); e.g., Figure 1C-E).

### Hierarchical clustering

To relate brain regions to phenomenological taxonomies, we filtered the region–category pairs with at least 10 total trials. For the remaining entries, we calculated the proportion of hits out of total trials and scaled the density by applying a logarithmic transformation of the total number of trials to account for the trial quantity. The resulting weighted density matrix was used as input to MATLAB’s clustergram function to perform hierarchical clustering, allowing us to group regions and categories with similar response profiles.

### Further refining motor findings

To refine the topography of motor findings, we performed a principal component analysis on the MNI152 Y and Z coordinates (i.e. rostral-caudal and dorsal-ventral coordinates). We then projected the discrete coordinates of each segmental motor component onto the first principal component (the dorsoventral axis).

Because stimulation trials involve electrode pairs spaced 1 cm apart that may lie in different regions, it can be unclear which electrode contributed to a functional response. To address this, we generated correlation matrices across all possible electrode–ROI localizations and calculated the probability of a positive response for each pair (Supplementary Figure 1).

### Exploring “canonical” regions

“Canonical” language areas—classically defined perisylvian regions such as Broca’s area in the inferior frontal gyrus and Wernicke’s area in the posterior superior temporal gyrus—have been shown in studies of predominantly tumor patients to carry a high probability and low inter-patient variability of language disruption when stimulated.[2] We tested whether similar distinctions could be drawn in an epilepsy population by assessing inter-patient probability and variability (Figure 4A-C). We used mouth motor localization as a reference. We aggregated all individual patient trial data into an HCP-MMP ROI table. To avoid bias from undersampled ROIs, we included only ROIs that had at least 10 total trials and at least one positive trial. We then bootstrapped the ROIs 1,000 times with replacement and computed the mean hit probability and variance per ROI, depicted as violin plots and color-coded ROI maps (Figure 4C).

### Defining visual and auditory naming topography

To test the lobe-specific spatial dispersion of auditory and visual naming in the temporal lobe and frontal lobe, we performed a kernel density analysis (Figure 4D-G). To accomplish this, we first performed a principal component analysis of the MNI152 Y and Z coordinates. Next, we created a contour plot of the coordinates in PC1/PC2 space and overlaid the resulting centroids and kernel distributions. A Hotelling’s test was used to determine whether the kernel distance between the two modalities was statistically significant.[29] Lastly, we evaluated the hit probabilities for each language modality across the gyri within each lobe.

### Predictors of anomia and speech arrest likelihood

We aimed to assess whether specific clinical variables predict a decreased likelihood of eliciting these phenomena during cortical stimulation. We conducted two separate generalized linear mixed-effects model (GLME) analyses: one to predict anomia elicited by stimulation in the temporal lobe, and the other to predict speech arrest elicited in the frontal lobe. Each model included an interaction between the presence of the lesion in the relevant lobe (temporal or frontal) and age at epilepsy onset, categorized as≤ 6 years versus *>* 6 years. Additional fixed effects included the duration of epilepsy, prior epilepsy surgery, race, and sex. A random intercept was specified for each patient to account for between-patient variability. The outcome variable was modeled using a binomial distribution with a logit link function, and statistical significance was defined as p *<* 0.05.

## Results

To construct an atlas that maps structure–function relationships, we first collected all DES results and categorized them into a hierarchical taxonomy (Figure 1A-B). We analyzed a large cohort from 2008-2023 encompassing 128 implants across 125 patients (3 re-implants; 83 left-sided implants; 32 right-sided; 10 bilateral; see Table 1 and Supplementary Table 2 for further information). The major categories in the taxonomy were language, sensory, and motor, with hierarchical branches specific to each major category (see Methods: Electrical stimulation protocol and Figure 1B). Of 3407 total stimulation trials, 298 trials were excluded due to high impedance (161 trials) or dural pain (137 trials) that precluded testing. Trials that elicited seizures (214 trials) or auras (35 trials) were not analyzed. Trials that elicited afterdischarges (736 trials) were used when clearing a contact but not in positive trials. There were 2124 remaining trials: 1130 positive trials with at least one phenomenon and 994 negative trials.

**Table 1.**
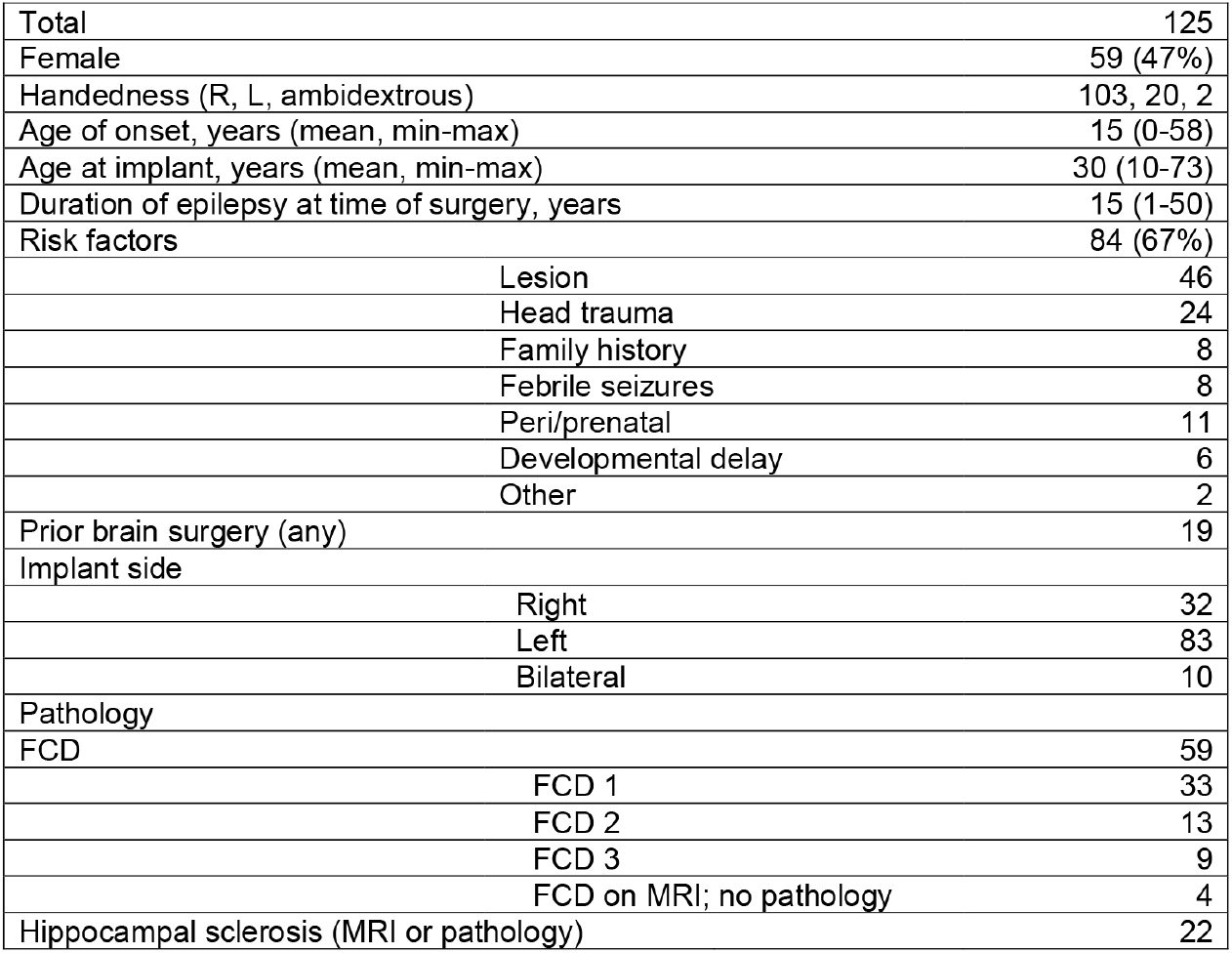
Patient characteristics.

### Distribution of major functional categories

We first mapped the cortical distribution and hit probability of primary language, sensory, and motor phenomena (Figure 1C-E). Language disruption (338 positive trials, 648 negative trials) was most frequently elicited in the posterior STG, parietal operculum, and frontal operculum (Figure 1C). Sensory phenomena (370 positive trials, 1554 negative trials) showed the highest concentration in the postcentral gyrus and supramarginal gyrus (Figure 1D). Motor responses (520 positive trials, 1572 negative trials) were predominantly localized to the precentral and postcentral gyri (Figure 1E). These distributions grossly reflected expected anatomical patterns, with each primary phenomenon (especially sensory and motor) localized to regions consistent with its functional domain. However, we elicited a large number of elementary motor phenomena (e.g., clonic or tonic movements) in contacts posterior to the central sulcus, extending into superior and inferior parietal cortex. This finding was seen even when both contacts in the stimulated pair were at a distance from the precentral gyrus (Supplementary Figure 1).

### Phenomena are segregated into phenomena- and ROI-specific hierarchical clusters

Given that most phenomena spanned multiple regions, we sought to identify prototypical profiles across the cortex without imposing assumptions about structure–function correspondence. We therefore applied hierarchical clustering as a data-driven method to organize widely distributed responses into meaningful functional and anatomical domains.(Methods: hierarchical clustering). This analysis identified three distinct clusters (Figure 2A-B).

**Fig. 2.**
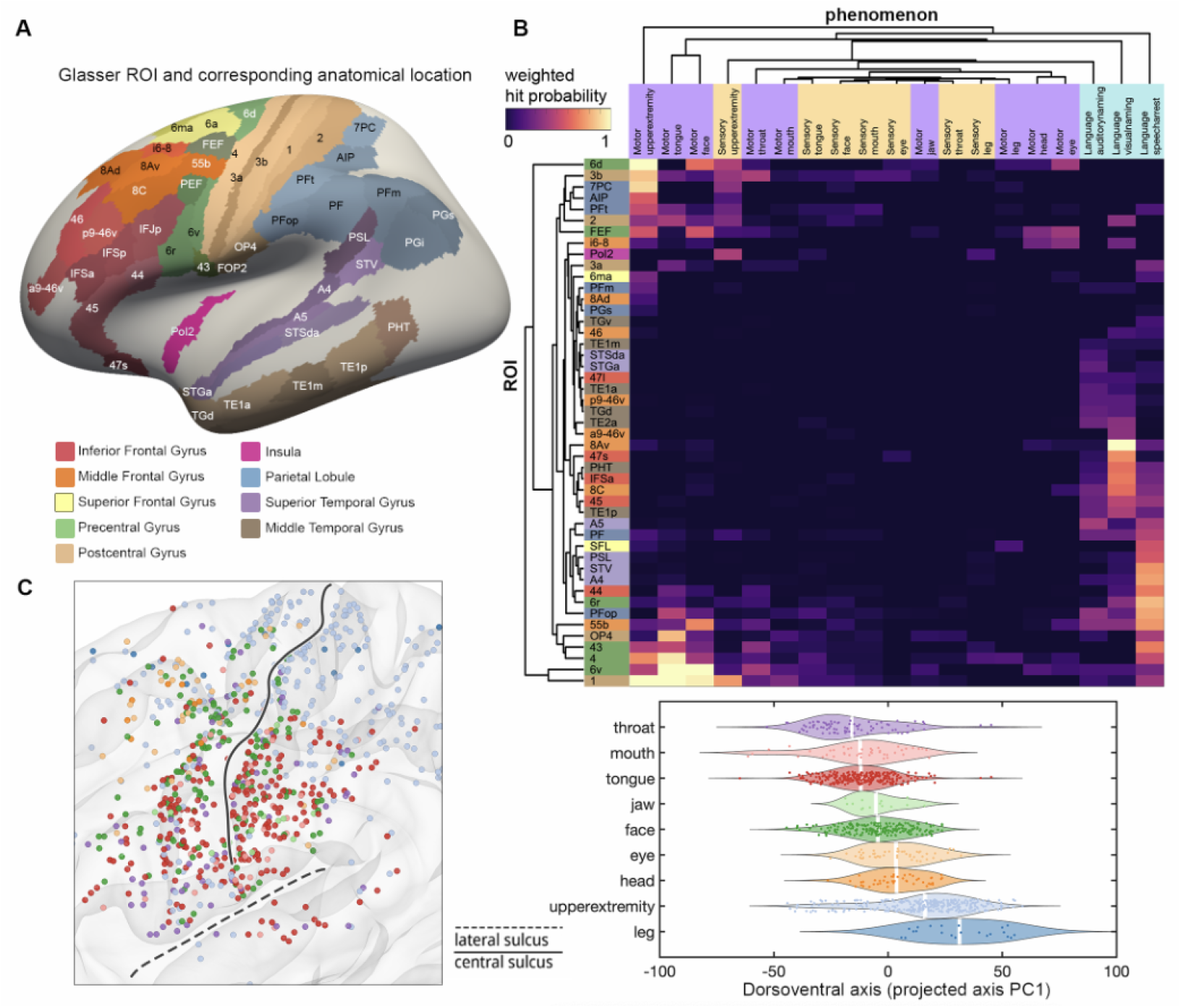
Hierarchical clustering of functional phenomena and motor somatotopy. (A) Lateral view of the Glasser parcellation in the left hemisphere colored by cortical region. (B) Hierarchical clustering of weighted hit probabilities across ROIs (rows) and elicited phenomena (columns). Color intensity indicates normalized likelihood of eliciting each phenomenon within a region. (C) Topographic organization of motor responses along precentral and postcentral gyri. Left: electrode locations color-coded by body regions, matching colors in the violin plot. Right: distribution of motor phenomena projected onto the dorsoventral axis (PC1) derived from electrode *MNI*_*y*_ and *MNI*_*z*_ coordinates. (D) Spatial distribution of throat and laryngeal responses in left (purple) and right (orange) hemispheres.

The first cluster consisted primarily of orofacial motor phenomena (Figure 2B, bottom left), including movements of the tongue, face, and throat, with strong representation in HCP-MMP areas 1, 4, 6v, and 43. The second cluster featured a sensorimotor-parietal network, encompassing both motor and sensory phenomena such as motor arm, motor face, and sensory hand (Figure 2B, top left). These were primarily located in more dorsal and posterior regions, including areas 6d, FEF, 2, and 3b, as well as parietal areas such as 7PC, AIP, and PFt. The third cluster captured language-related phenomena (Figure 2B, right), including auditory and visual naming and speech arrest. These were grouped based on their shared functional identity rather than spatial proximity, and were distributed across frontal, temporal, and parietal cortices. Notably, this cluster was centered on areas 8Av, 44, PFop, 6r, A4, and STV, concordant with areas previously identified as involved in cortical language networks. We also confirmed the dorsoventral somatotopic organization of motor phenomena along the central sulcus, with leg/foot motor phenomena localized most dorsally, followed by arm/hand/finger phenomena, and finally eye and head representations most ventrally and anteriorly (Figure 2C).

### Spatial probability and variability of language disruption

To systematically resolve heterogeneous and spatially dispersed language findings, we applied a multi-level analytic framework that unifies discrete coordinate analyses (Figure 3A), probability distributions (Figure 3B), ROI-based bootstrapping (Figure 3C), and kernel density estimation (Figure 3D–G). We identified 338 trials with positive language interference and 648 trials in which adequate stimulation failed to disrupt language. Of the positive trials, 335 were in the left hemisphere and 3 trials (from two patients) were in the right hemisphere.

**Fig. 3.**
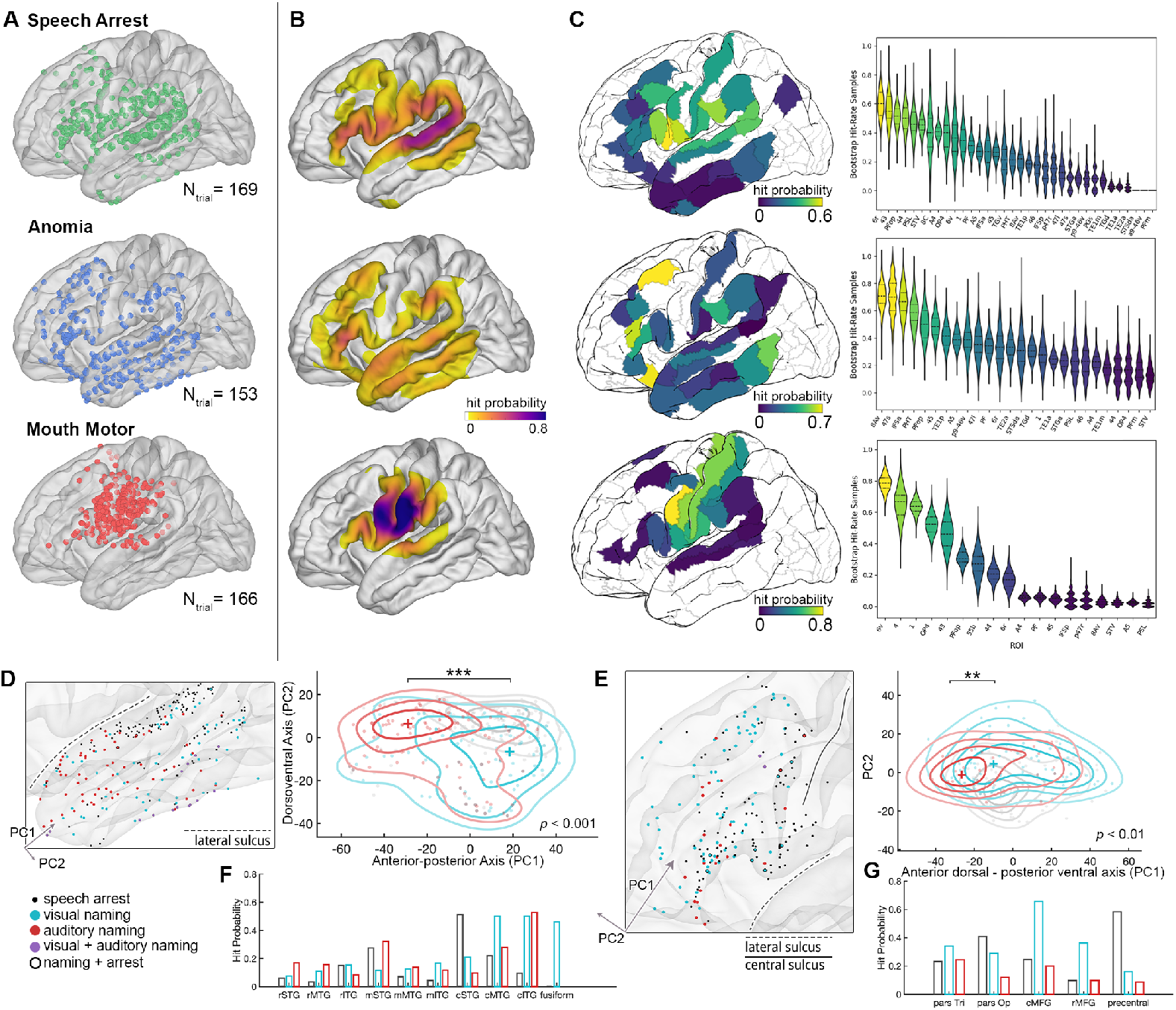
Estimates of hit probability and variability for language phenomena. (A) Raw electrode distributions for three language-related phenomena: anomia, speech arrest, and mouth motor disruption. Each dot represents a single electrode site. (B) Cortical hit probabilities for each phenomenon. (C) Left: cortical map showing the sorted mean probability across Glasser ROIs. Right: violin plots sorted by mean hit probability, showing bootstrapped hit rate distributions per ROI; color on both the cortex and violin plots represents the ROI’s mean hit probability. (D) and (E): Left: zoomed-in view of language disruption in temporal lobe and frontal lobe, respectively, with each electrode color-coded by language disruption and their co-occurrence: visual naming (blue), auditory naming (red), or speech arrest (black). Right: kernel distribution of each disruption, significance of kernel distance tested by Hotelling’s test. (F) and (G) breakdown of hit probability across anatomical regions.

**Fig. 4.**
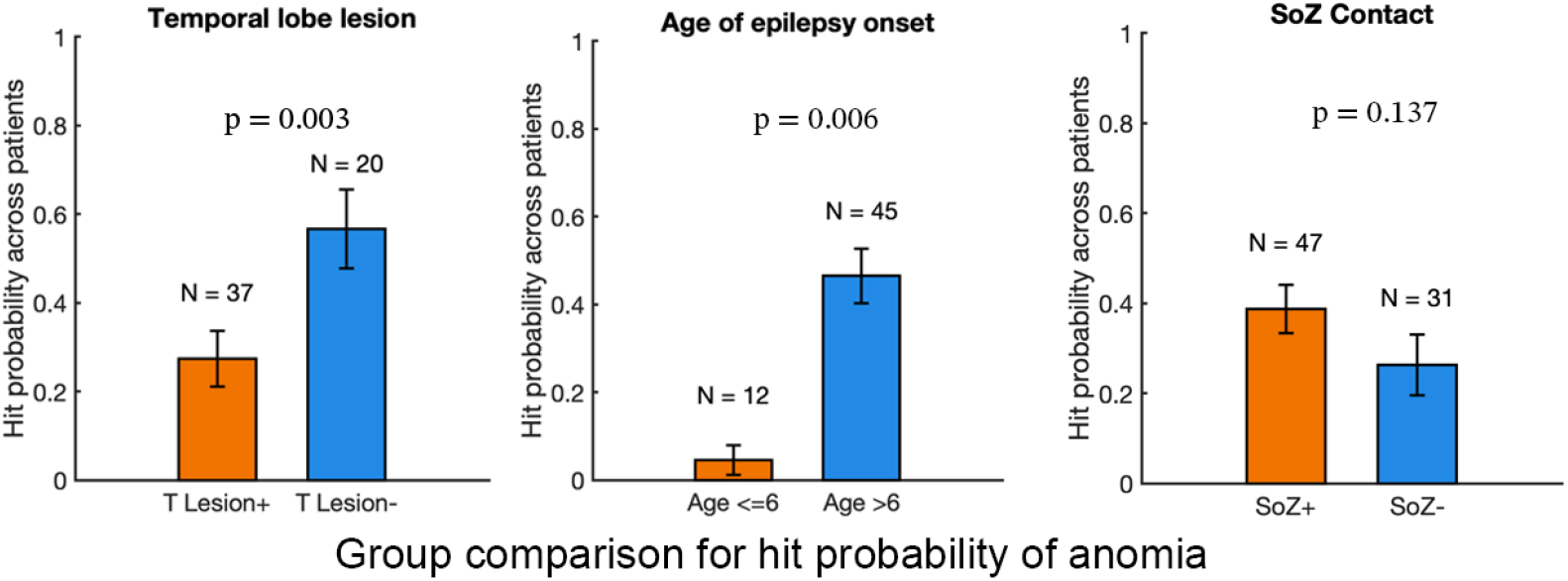
Group-level predictors of anomia. (A) Patients with temporal lobe lesions (T Lesion+) exhibited significantly lower anomia hit probability compared to those without temporal lobe lesions (T Lesion–; *p* = 0.003). Analysis was restricted to stimulation delivered to temporal lobe electrodes. (B) Patients with epilepsy onset age ≤ 6 years showed significantly reduced anomia hit probability compared to those with later onset (*>* 6 years; *p* = 0.006). (C) No significant difference in anomia hit probability was observed between electrodes located within the seizure onset zone (SOZ+) versus those outside (SOZ–). All comparisons were performed at the patient level using linear mixed-effects models. Panels B and C include stimulation from electrodes across the entire brain.)

Distinct spatial patterns emerge for speech arrest vs. anomia. Speech arrest responses were concentrated around perisylvian regions, particularly the inferior frontal and peri-Rolandic cortices (figure 3A, top). High hit probabilities were observed in areas PFop, 6r, and 43/44 (Figure 3B and 3C, top). Anomia trials (visual naming: 99 hits; auditory naming: 58 hits) were more broadly distributed, encompassing the pars triangularis, caudal MFG, and lateral temporal cortex. The highest probabilities were found in areas 8Av, 47s, and IFSa (Figure 3A-C, middle).

Mouth motor interference was included in the analysis given its role in the language testing battery and its impact on subsequent assessment. Hits were predominantly located in the pre- and postcentral gyri, with occasional extension into prefrontal cortex. Areas 6v, 4, and 1 exhibited high hit probabilities and low variability (Figure 3A-C, bottom). This contrasts with the higher variability seen in speech arrest and anomia areas.

### Auditory and visual naming topography

Prior studies suggests that visual and auditory anomia and speech arrest follow consistent topographies within the temporal and frontal lobes.[11, 30] In our cohort, auditory and visual naming disruptions demonstrated an anterior-posterior dissociation within the temporal lobe, with auditory naming responses located anteriorly while visual naming responses clustered posteriorly. Despite overlap, the kernel density distributions of these two modalities were significantly different (Hotelling’s test, *p <*1e-5; Figure 3D, right; Figure 3F).

In the frontal lobe, auditory and visual naming hits organized along a combined anterior/dorsal to posterior/ventral axis (Figure 3E and 3G). Auditory naming hits were more likely to be localized in antero-ventral regions, whereas visual naming hits occurred more frequently in postero-dorsal sites. This spatial separation was again significant by kernel density comparison (Hotelling’s test, *p* = 0.0094; Figure 4G).

### Predictors of visual or auditory naming disruption

We tested the hypothesis that certain clinical variables predict a lower probability of naming disruption. In a generalized linear mixed effects model that included age of epilepsy onset ≤ 6 and presence of a temporal lobe lesion as well as their interaction and accounted for within-subject repeated measures, both temporal lobe lesion and age of onset ≤ 6 predicted a decreased likelihood of anomia hits in the temporal lobe (temporal lobe lesion: 17/49 vs. 15/28, *p* = 0.003; age of onset ≤ 6: 2/14 vs. 30/63, *p* = 0.006; Figure 4A-B). When controlling for lesion and age of onset, the electrode being within the seizure onset zone did not independently predict a naming hit (linear mixed effect model, *p* = 0.137; Figure 4C).

### Higher-order and experiential phenomena

We observed a broad range of higher-order and experiential phenomena which do not fit neatly into the above rudimentary clusters. To spatially characterize these phenomena, we mapped the raw distributions drawing from 45 patients and 137 stimulation trials (Figure 5A). Complex sensory hallucinations followed modality-specific topographies: auditory percepts were concentrated in the posterior STG, particularly posterior regions, while visual hallucinations were concentrated in the occipital lobe. Vertiginous sensations were evoked primarily from the insula, posterior temporal, and parietal cortices. We highlight two reproducible experiential phenomena (Figure 5B). A feeling of presence was reported by seven patients in eight trials. This percept was the sensation of someone standing behind or next to the patient, with or without an auditory component (e.g., hearing speech or counting) and elicited by stimulation of posterior STG, MTG, and supramarginal gyrus. Six patients reported forced thoughts or memories (e.g., thoughts of a specific person, game or television commercial) with stimulation of electrode pairs in the frontal and temporal lobes; the majority of these were in rostral MFG (Figure 5B).

**Fig. 5.**
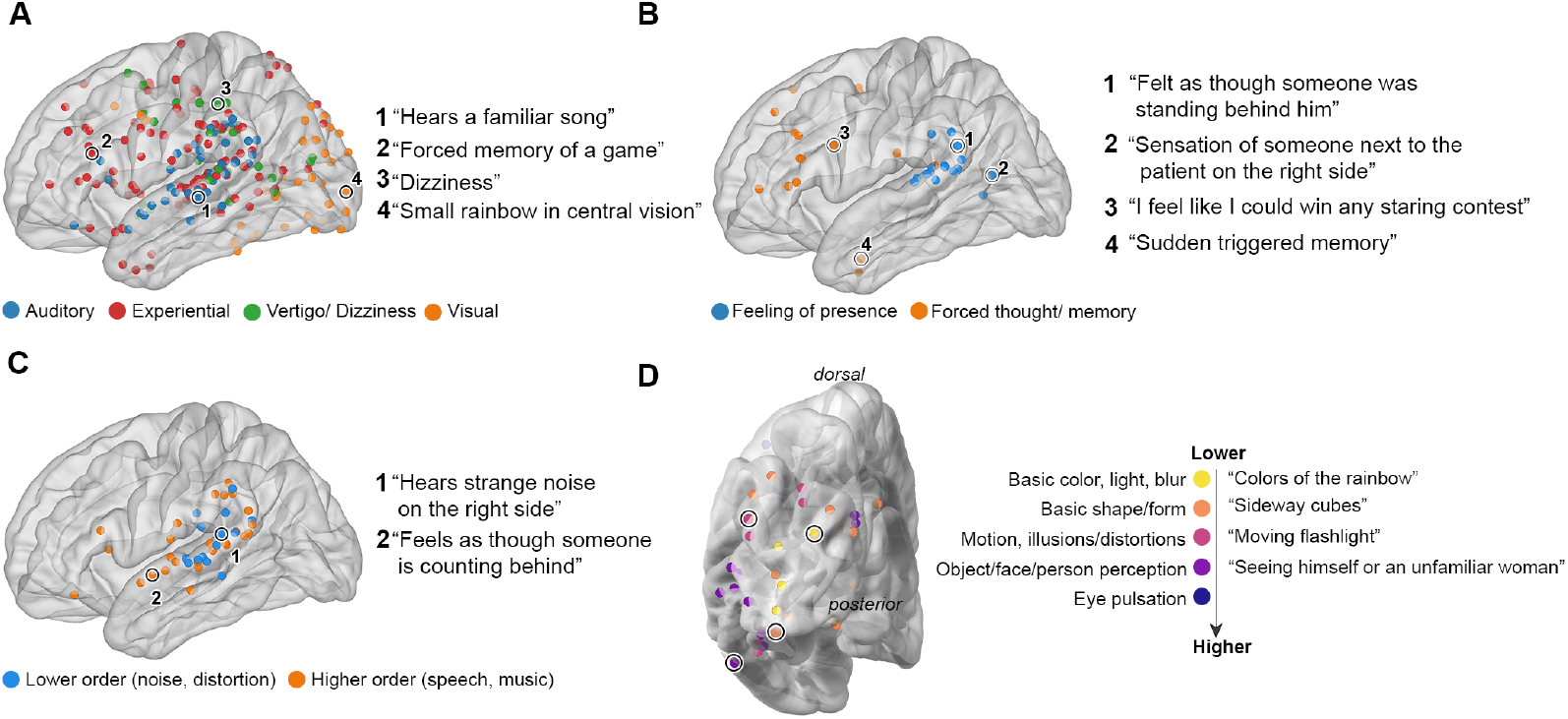
Spatial distribution of non-somatosensory and experiential phenomena. (A) Cortical distribution of reported percepts, color-coded by category: auditory (blue), experiential (red), vertigo/dizziness (green), and visual (orange). Numbered labels correspond to representative examples. (B) Mapped stimulation sites for two reproducible experiential phenomena: feeling of presence (blue) and forced thoughts or memories (orange). (C) Auditory hallucinations divided into lowerorder (blue; e.g., noise, distortion) and higher-order (orange; e.g., speech, music) percepts. (D) Spatial gradient of visual hallucinations mapped onto occipital cortex, progressing from low-level features (e.g., color, shape) to high-order percepts (e.g., object and face imagery), shown with color-coded categories.

We categorized auditory hallucinations into lower-order (e.g., buzzing, static) and higher-order (e.g., speech, music) phenomena (Figure 5C, 14 patients, 27 trials). Lower-order responses were clustered in posterior STG and supramarginal regions, while higher-order responses spanned mid-to-anterior STG and occasionally extended into inferior frontal areas. These higher-order responses were often multisensory, combining auditory with emotional or visual features.

Visual hallucination data from 30 stimulation trials across eight patients, with one patient contributing 21 trials from extensive occipital lobe coverage (Figure 5D). There was a spatial progression of visual phenomena consistent with established visual processing hierarchies. Stimulation of early visual areas (e.g., V1/V2) elicited simple percepts (e.g., diffuse light, color, or basic shapes) whereas stimulation of more anterior occipital regions evoked increasingly complex visual experiences (e.g., including motion, object, and face-related imagery).

## Discussion

We present a probabilistic sensory, motor, and language atlas based on ≥ 2,000 extraoperative DES trials across 125 patients, supporting that cortical functional organization is better characterized as a probabilistic continuum rather than discrete “canonical” regions in an epilepsy population. This contrasts with intraoperative studies of patients primarily with tumors with or without epilepsy, who can have more tightly clustered “canonical” regions.[2, 6] Our results underscore the importance of individualized and extensive testing for functional brain areas in people with epilepsy.

### The probability of language disruption exists on a continuum

Our findings reveal that language-disruption probabilities vary smoothly across the cortex without sharp boundaries (Figure 3). In contrast, prior DES studies reported disruption maps by calculating the density of positive sites or by reporting the proportion of patients with at least one positive response [3, 5, 6]. Because these approaches do not normalize by the total number of stimulations delivered at each location (denominator), the “probabilistic” conclusions are subject to sampling bias and overrepresented ROIs; without denominators, a high density region may simply reflect frequent testing rather than probability. Moreover, intraoperative DES typically samples sparsely outside peri-sylvian cortices, limiting the ability of prior studies to derive reliable ROI-based probability or density estimates. [3, 6].

Our study addresses these concerns by calculating the probability of disruption at millimeter and ROI levels, using a true denominator that includes positive and negative stimulation trials (i.e., cleared sites that passed maximal stimulation without eliciting a response) across broad cortical coverage. The speech arrest probability map shows the highest mean probability in ventral precentral and subcentral gyrus (area 6r and 43) and pars opercularis (Figure 3C, top), in line with results in hit density [2, 6]. Further, anomia shows its highest probability spanning IFG and MFG (8Av, 47s, IFSa, and area 45), and posterior STG and MTG (area PHT, PFop, and TE1p) (Figure 3C, middle). With the fine parcellation, we showed, for the first time, a graded probabilistic distribution without sharp drop-offs across the cortex. These findings provide near-causal evidence for theoretical frameworks describing the distributed organization of speech and language function [15, 25, 31].

Although significant inter-patient variability was seen, we did identify certain predictors of language reorganization. In our large cohort, a temporal lobe lesion was associated with a lower likelihood of visual/auditory naming disruption elicited in the temporal cortex (Figure 5A), but not in the frontal lobe, consistent with prior intraoperative and extraoperative studies. [10, 13] Similarly, seizure onset before age 6 predicted a lower likelihood of anomia in the frontal and temporal lobes (Figure 5A). However, when controlling for lesion status and age of epilepsy onset, the seizure onset in a language region did not predict anomia probability.

Our study also identifies MFG as a robust site of language disruption (Figure 3). In a detailed breakdown of all language disruptions in 15 patients (Supplementary Figure 2A, left), naming and speech arrest span MFG, consistent with prior reports from iEEG of its involvement in speech planning and production.[32–35] A finer parcellation with co-stimulation distribution reveals areas 8Av and 8C the highest within-pair percentage (28/36 pairs in 8Av, and 8/8 in 8C), but language disruption does not occur when both stimulated contacts were in area 55b (Supplementary Figure 2B); its involvement only occurs with co-stimulation of 8Av or 8C electrodes (both triggered 100% hit). In contrast, 55b shows stronger involvement in motor phenomena (8/24 within-pair positive hits, and near 100% hit rate when co-stimulated with areas 2, 3b, 4, 6d, and FEF), suggesting a role in speech motor production rather than non-motor language processes (see Supplementary Figure 1, top)[36, 37]. Together, these findings provide causal evidence for 8Av/8C in naming and speech arrest, while refining the role of 55b as a node supporting speech motor production.[15, 38]

Our results also provide systematic spatial segregation of the neural substrate in auditory and visual naming.Figure 3D and E Auditory naming localization is of special interest given its role in predicting risks of post-operative word retrieval difficulty that cannot be detected by visual naming alone. ([30, 39, 40]) DES studies have rarely examined this task intraoperatively, with reported effects largely confined to the temporal lobe.[30, 39, 41] We showed that in the prefrontal cortex, auditory naming engaged more anteroventrally, whereas visual naming engaged more posterodorsally. These findings provide causal evidence for a modality-specific gradient consistent with recent ECoG studies (Figure 3D).[33, 42] These results support a distributed, modality-sensitive organization of naming beyond classical temporal loci.

### Motor and sensory findings

Motor responses in our cohort were not confined to the precentral gyrus but extended posteriorly into postcentral and parietal cortices. Although classically considered sensory cortex, Penfield first noted motor responses posterior to the central sulcus[7] and subsequent reports have confirmed that stimulation of postcentral gyrus can elicit positive or negative motor phenomena.[43] More recent stereo EEG and intraoperative DES studies have demonstrated motor or multimodal responses in supramarginal and superior parietal lobules, suggesting these regions can play a role in elementary motor control as well as higher-order motor control and sensorimotor integration.[44, 45] Our findings reinforce and extend these observations in a large extraoperative DES cohort. We found robust and reproducible motor effects even when both stimulating contacts were located at a distance from the precentral gyrus and provide a probabilistic map of motor responses in the parietal cortex (Supplementary Figure 1). Clinically, these findings argue for broad functional DES mapping when resections approach parietal association cortex.

Although higher-order experiential phenomena have been described at length from neuroimaging and seizure semiology studies, reports of DES-induced phenomena are rare. We contribute several cases of DES-induced feeling of presence and forced thinking to the literature (Figure 5). In our cohort we had seven patients who reported a feeling of presence during electrical stimulation of the temporoparietal junction. Feelings of presence are well-described as a result of numerous neurological and psychiatric diseases.[46] It is also reported as a seizure semiology where it tends to localize to the frontoparietal, temporoparietal, and insular cortices.[47] It is thought to be caused by a disruption of sensory integration pathways important for ones spatial sense of their own body.[46] Few cases have been reported induced by DES.[48, 49] To our knowledge, we offer the largest collection of DES-induced feelings of presence to date.

We induced forced thoughts or intrusive memories, which has only previously been reported in small case series (including a prior report of three patients from this dataset[50]). As a seizure semiology, forced thought or hypercognitive seizures[51] usually localizes to the frontal lobe.[52–55] However, the concept of forced thinking and forced memories can be conflated by patients and clinician interpretations;[51] temporal lobe seizures and stimulation near the arcuate fasciculus[51] can also evoke a similar phenomenon.[56]

### Limitations

Our study has notable limitations. First, although this was a large cohort with bilateral hemispheric representation, most patients were right-handed. Although most lefthanded and anomalous dominance patients had left language representation, we had insufficient left-handed or anomalous dominance patients for subgroup exploration. Second, patients did not have complete mapping of all implanted electrodes, leading to variability in inter-patient cortical representation. This limitation is partly overcome by using an ROI-based approach and bootstrapping from a large dataset. Lastly, although phenomena could be seen at a distance from canonical sites (e.g., positive motor phenomena in the superior parietal lobule), it is not known whether this reflects propagated synaptic activation or whether a cortical resection in these “non-canonical” locations would result in a similar deficit as it would from a resection of a “canonical” location such as primary motor cortex.

## Conclusion

Our analyses provide a practical atlas based on extraoperative DES of motor, sensory, and language functions, mapped in both MNI coordinates and Glasser’s functional parcellation. We generated probabilistic maps of motor, sensory, and language responses to help identify and delineate cortical regions of critical function. While the results align with known patterns of cortical organization, the marked inter-subject variability underscores the importance of individualized functional mapping in pre-surgical planning.

## Acknowledgments

This work was supported by National Institute of Health grants R01NS109367, R01NS115929, and R01DC018805 (A.F.). A.J.M. is supported by the American Epilepsy Foundation Research & Training Fellowship for Clinicians.

## Data and Code availability

The data set generated during the current study will be made available from the authors upon request and documentation is provided that the data will be strictly used for research purposes and will comply with the terms of our study IRB. The code is available upon publication at https://github.com/flinkerlab/.

## Declarations

The authors declare that they have no competing interests.

## Supplementary Information

**Fig. S1.**
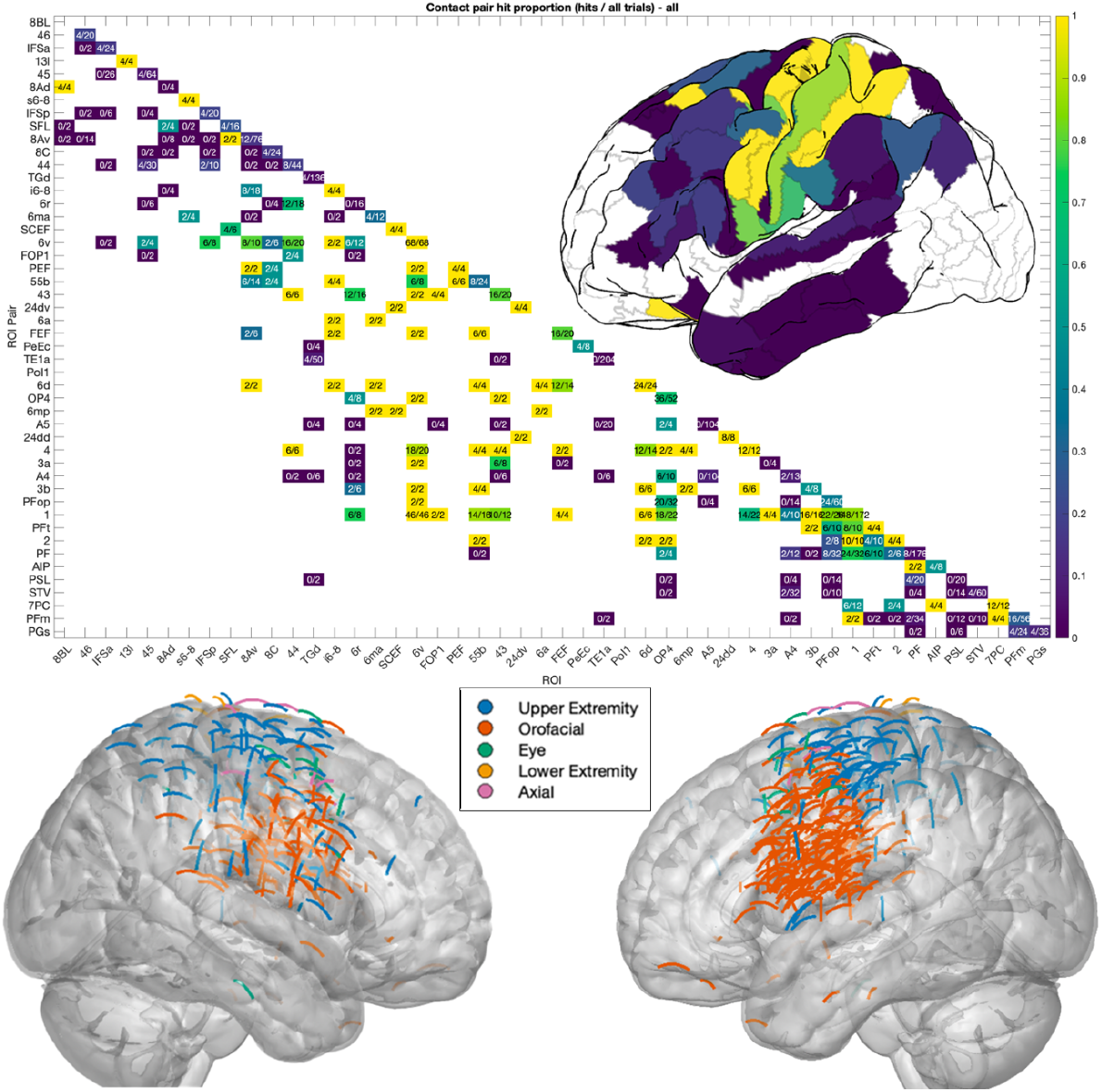
Proportion of positive trials by contact pair location. A: Matrix depicting the proportion of positive motor trials based on the ROI of each contact in the stimulated pair. The diagonal represents trials where both contacts in the pair were within the same ROI. Inset: Illustration of the hit proportions from the diagonal of the matrix. B: Positive motor trials with each arc representing the pair of contacts that contributed to a positive trial, grouped into five major categories. All panels illustrate that motor phenomena can be elicited at a distance posterior to the central sulcus.

**Fig. S2.**
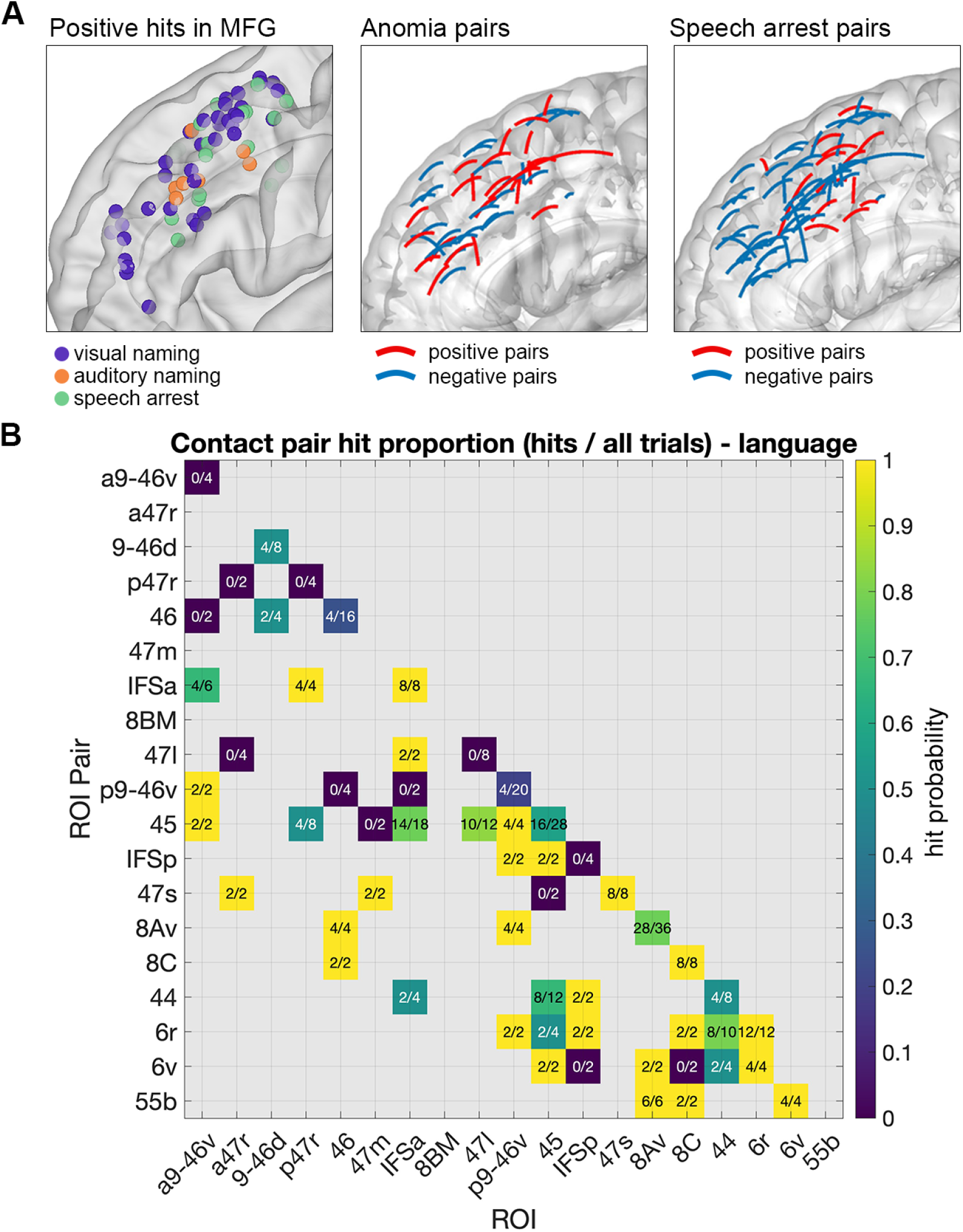
Details of phenomena evoked by DES of MFG. (A) Left: all positive phenomena in MFG. Middle: contact pair location for anomia. Positive trials are in red and negative trials are in blue. Right: contact pair location for speech arrest. (B) Matrix depicting the proportion of positive language trials based on the ROI of each contact in the stimulated pair in prefrontal regions. The diagonal represents trials where both contacts in the pair were within the same ROI.

